# Vaccine-mediated protection against merbecovirus and sarbecovirus challenge in mice

**DOI:** 10.1101/2023.05.22.540829

**Authors:** David R. Martinez, Alexandra Schaefer, Tyler D. Gavitt, Michael L. Mallory, Esther Lee, Nicholas J. Catanzaro, Haiyan Chen, Kendra Gully, Trevor Scobey, Pooja Korategere, Alecia Brown, Lena Smith, Rob Parks, Maggie Barr, Amanda Newman, Cindy Bowman, John M. Powers, Katayoun Mansouri, Robert J. Edwards, Ralph S. Baric, Barton F. Haynes, Kevin O. Saunders

## Abstract

The emergence of three distinct highly pathogenic human coronaviruses – SARS-CoV in 2003, MERS-CoV in 2012, and SARS-CoV-2 in 2019 – underlines the need to develop broadly active vaccines against the *Merbecovirus* and *Sarbecovirus* betacoronavirus subgenera. While SARS-CoV-2 vaccines are highly protective against severe COVID-19 disease, they do not protect against other sarbecoviruses or merbecoviruses. Here, we vaccinate mice with a trivalent sortase-conjugate nanoparticle (scNP) vaccine containing the SARS-CoV-2, RsSHC014, and MERS-CoV receptor binding domains (RBDs), which elicited live-virus neutralizing antibody responses and broad protection. Specifically, a monovalent SARS-CoV-2 RBD scNP vaccine only protected against sarbecovirus challenge, whereas the trivalent RBD scNP vaccine protected against both merbecovirus and sarbecovirus challenge in highly pathogenic and lethal mouse models. Moreover, the trivalent RBD scNP elicited serum neutralizing antibodies against SARS-CoV, MERS-CoV and SARS-CoV-2 BA.1 live viruses. Our findings show that a trivalent RBD nanoparticle vaccine displaying merbecovirus and sarbecovirus immunogens elicits immunity that broadly protects mice against disease. This study demonstrates proof-of-concept for a single pan-betacoronavirus vaccine to protect against three highly pathogenic human coronaviruses spanning two betacoronavirus subgenera.

## INTRODUCTION

The emergence of SARS-CoV in 2003, MERS-CoV in 2012, and SARS-CoV-2 in 2019 into naïve human populations underlines the spillover potential of coronaviruses. SARS-CoV-2 causes coronavirus disease of 2019 (COVID-19) ^1^. The COVID-19 pandemic has had a devastating impact on human health and the world economy. SARS-CoV, SARS-CoV-2, and several zoonotic, pre-emergent SARS- and SARS2-related bat coronaviruses belong to the *Betacoronavirus* genus and *Sarbecovirus* subgenus and are classified as Group 2b coronaviruses ^2–4^. Similarly, MERS-CoV and MERS-related bat zoonotic viruses also belong to the *Betacoronavirus* genus and *Merbecovirus* subgenus and are classified as Group 2c coronaviruses ^2, 3^. Given that in the last two decades, one merbecovirus and two sarbecoviruses have emerged into humans, the development of countermeasures against these important groups of viruses – including universal coronavirus vaccines –is a global health priority.

Several pan-betacoronavirus vaccine approaches have shown early promise in animal models ^5–8^. Sortase-conjugated ferritin nanoparticles (scNPs) bearing the SARS-CoV-2 receptor binding domain (RBD) elicited neutralizing antibodies against bat SARS-related viruses and protected non-human primates (NHP) against SARS-CoV-2 challenge ^9^. Moreover, monovalent SARS-CoV-2 RBD scNP vaccines elicit neutralizing antibodies against all tested SARS-CoV-2 variants including D614G, Beta, Delta, Omicron BA.1, BA.2, BA.2.12.1, and BA.4/BA.5 ^10^.

Similar approaches with RBD nanoparticle vaccines also protect against sarbecovirus challenge in mice ^8^. Chimeric spike antigens delivered as multiplexed mRNA-LNP vaccines similarly protected mice from genetically divergent bat zoonotic SARS-related viruses and SARS-CoV-2 variants ^6^. Therefore, multiple vaccine designs and modalities have protected against heterologous sarbecovirus challenge in animal models. Importantly, humans infected with SARS-CoV 2003 and/or SARS-CoV-2 generate neutralizing monoclonal antibodies capable of neutralizing SARS-related zoonotic viruses and SARS-CoV-2 variants ^11–15^. These human monoclonal antibodies protected mice and monkeys from sarbecovirus infection ^11, 16^. These studies indicated that elicitation of protective neutralizing antibody responses against sarbecoviruses is achievable.

Despite demonstrating proof-of-principle that vaccines can elicit broad immunity against genetically divergent sarbecoviruses ^5–8, 17^, no study to date has demonstrated vaccine-mediated protection against both sarbecovirus and merbecovirus betacoronaviruses. While stem-helix antibodies isolated from humans can protect against Group 2b SARS-CoV and SARS-CoV-2 as well as Group 2c MERS-CoV in highly pathogenic mouse models ^18^, current vaccination strategies do not reproducibly induce immunity targeting these conserved S2 epitopes. Therefore, alternative vaccination strategies that effectively target sarbecoviruses and merbecoviruses are needed.

SARS-CoV-2 spike mRNA vaccines do not protect mice against challenge with genetically divergent zoonotic SARS-related viruses and SARS-CoV ^6^. This suggests that currently used SARS-CoV-2 mRNA spike vaccines are unlikely to strongly protect against future SARS-related, SARS-CoV-2-related zoonotic viruses, or highly evolved SARS-CoV-2 variants of concern that could emerge in the future ^19, 20^. We therefore developed a trivalent RBD vaccine composed of sarbecovirus and merbecovirus RBDs from zoonotic pre-emergent, human epidemic, and pandemic coronaviruses. In this study, we evaluated the immunogenicity and protective efficacy against SARS-CoV and MERS-CoV in mice. We show that a monovalent SARS-CoV-2 RBD nanoparticle can protect against heterologous sarbecovirus challenge but does not protect against merbecovirus challenge. Conversely, the trivalent RBD scNP generates neutralizing antibodies and prevents severe sarbecovirus disease and merbecovirus infections. This study demonstrates proof-of-concept in an *in vivo* challenge setting that a vaccine that protects against merbecoviruses and sarbecoviruses is an achievable goal.

## RESULTS

### Generation and validation of trivalent RBD ferritin nanoparticle vaccine

We previously reported that a ferritin scNP monovalent SARS-CoV-2 RBD vaccine elicited broadly neutralizing antibodies against bat zoonotic pre-emergent betacoronaviruses, SARS-CoV, and SARS-CoV-2 variants in non-human primates ^7, 16^. To broaden the response of this SARS-CoV-2 RBD vaccine, we sought to generate a vaccine that increased the immunogenicity against the high risk *Merbecovirus* (also called Group 2c coronavirus) subgenus of betacoronaviruses, which includes MERS-CoV ^2^. We designed a trivalent sortase-A-conjugated 24-mer ferritin nanoparticle (scNP) vaccine displaying SARS-CoV-2 RBD, SARS-related bat RsSHC014 RBD, and MERS-CoV EMC RBD (Figure 1A and S1A) ^10^. Equimolar ratios of each RBD were conjugated to 24 acceptor sites on the 24-mer ferritin nanoparticle. In addition to the sarbecovirus SARS-CoV-2 RBD, RsSHC014 RBD was chosen for inclusion because it is a pre-emergent ACE2-binding sarbecovirus ^19^ to which the SARS-CoV-2 RBD nanoparticle generated only low levels of neutralizing antibodies ^7, 16^. We used negative stain electron microscopy (NSEM) to visualize the sortase A conjugated trivalent vaccines and demonstrated successful RBD conjugation (Figure 1B and S1B). The trivalent RBD scNP recapitulated the stability of the individual RBDs indicating the conjugation reaction had no deleterious effects on RBD folding or stability (Figure S1C and D).

**Figure 1.**
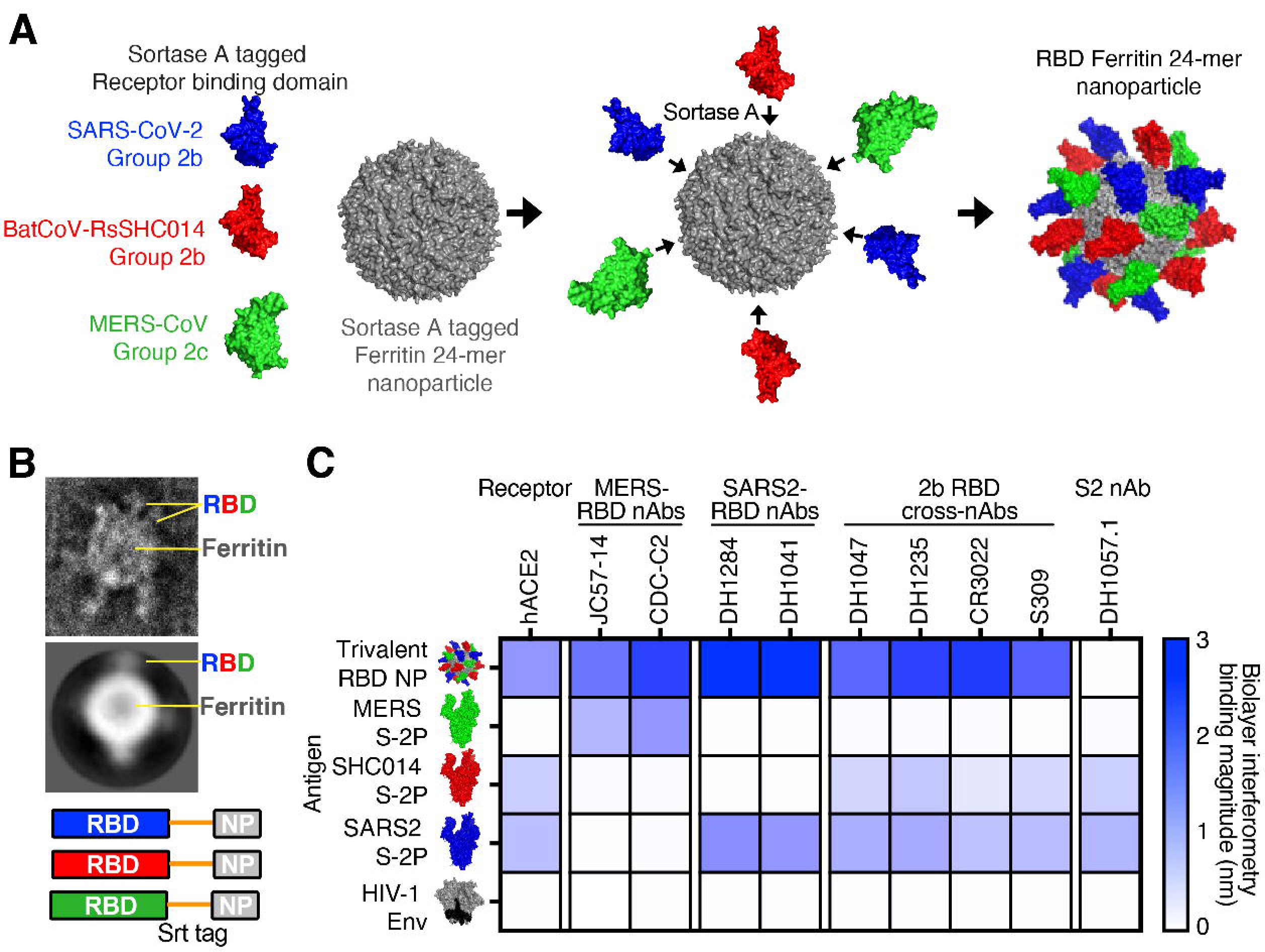
Design and characterization of trivalent RBD scNP vaccines. (**A**) Ferritin nanoparticles were conjugated with sortase A tagged Group 2b SARS-CoV-2 RBD, Group 2b RsSHC014 RBD, and Group 2c MERS-CoV RBD. (**B**) Visualization of trivalent scNP was performed via negative stain electron microscopy. (**C**) Validation of trivalent scNP vaccine by Biolayer Interferometry. Trivalent RBD scNP antigenicity was done by assessing binding of the trivalent vaccine and various Group 2b and Group 2c spikes to human ACE2, MERS-CoV RBD mAbs, SARS-CoV-2 RBD mAbs, Group 2b cross-reactive RBD mAbs, and an S2 mAb. HIV-1 envelope was included as a negative control antigen.

To validate the efficient conjugation of SARS-CoV-2/RsSHC014/MERS-CoV RBDs as a trivalent vaccine, we also performed biolayer interferometry (BLI) binding analyses with human monoclonal antibodies that recognize Group 2b and Group 2c coronavirus spike epitopes and human ACE2. MERS-CoV RBD-specific mAbs JC57-14 and CDC-C2 only recognized MERS-CoV spike and the trivalent RBD vaccine (Figure 1C). Similarly, SARS-CoV-2 RBD-specific mAbs DH1284 and DH1041 bound only to SARS-CoV-2 spike and the trivalent RBD vaccine (Figure 1C). Group 2b RBD cross-reactive mAbs DH1047, DH1235, CR3022, and S309 bound to SARS-CoV-2 spike, RsSHC014 spike, and the trivalent RBD vaccine with highest magnitude but not to MERS-CoV spike or HIV Env. Finally, the negative control stem-helix mAb DH1057.1 bound to RsSHC014 spike and SARS-CoV-2 spike but not to the trivalent RBD vaccine, MERS-CoV spike, or HIV Env. Overall, the trivalent RBD nanoparticle bound to all the various Group 2b and 2c RBD antibodies, whereas no one spike protein recapitulated this breadth of reactivity. These BLI binding analyses suggest that the trivalent SARS-CoV-2/RsSHC014/MERS-CoV RBD scNP vaccine was efficiently conjugated and the RBD immunogens are properly recognized by various Group 2b and Group 2c-reactive monoclonal antibodies.

### Immunogenicity of monovalent versus trivalent scNP vaccines in mice

To compare the immunogenicity of the monovalent versus the trivalent RBD scNP vaccines, we vaccinated aged BALB/c two times four weeks apart (Figure 2A). The Toll-like receptor 4 agonist glucopyranosyl lipid adjuvant-stable emulsion (GLA-SE) was used as the adjuvant for both vaccine groups and adjuvant-only controls (Figure 2A) ^21^. We immunized mice with 10μg of the monovalent SARS-CoV-2 RBD scNP vaccine and also 10μg of the trivalent SARS-CoV-2/RsSHC014/MERS-CoV RBD scNP vaccine adjuvanted with 5μg of the GLA-SE adjuvant. We measured serum binding IgG antibodies against sarbecovirus and merbecovirus spike ectodomain matching the RBDs present in the vaccine and SARS-CoV spike, which was not in the vaccine. In mice vaccinated twice with the SARS-CoV-2/RsSHC014/MERS-CoV trivalent RBD scNP vaccine, high titers of spike binding IgG antibodies against human outbreak SARS-CoV Tor2 isolate (Figure 2B), bat pre-emergent RsSHC014 (Figure 2C), the SARS-CoV-2 Wuhan-1 outbreak isolate (Figure 2D), and the MERS-CoV EMC isolate (Figure 2E) were observed. In agreement with the IgG binding to various Group 2b and Group 2c spikes, we also observed serum antibody blocking of hACE2 binding to SARS-CoV-2 Spike and hDPP4 binding to MERS-CoV in trivalent RBD scNP vaccinated mice (Figure 2F and 2G). Only the trivalent scNP vaccine elicited robust serum antibody responses capable of blocking hDPP4 (Figure 2G). The monovalent SARS-CoV-2 RBD scNP vaccine also elicited high titers of binding IgG antibodies two weeks post boost against the three sarbecovirus spikes SARS-CoV Tor2 isolate, RsSHC014, SARS-CoV-2 Wuhan-1 isolate and human ACE2-blocking antibodies (Figure 2B-D, 2F). However, immunization with SARS-CoV-2 RBD scNP did not elicit binding IgG to MERS-CoV spike or DPP4-blocking antibodies (Figure 2E 2G). These data indicated that the monovalent SARS-CoV-2 RBD scNP vaccine elicited cross-reactive binding and serum-blocking antibodies to Group 2b but not Group 2c coronaviruses. Thus, the trivalent RBD scNP vaccine improved antibody responses compared to the monovalent SARS-CoV-2 RBD scNP by eliciting serum antibodies to spikes from all three highly pathogenic human betacoronaviruses and a pre-emergent bat coronavirus.

**Fig. 2.**
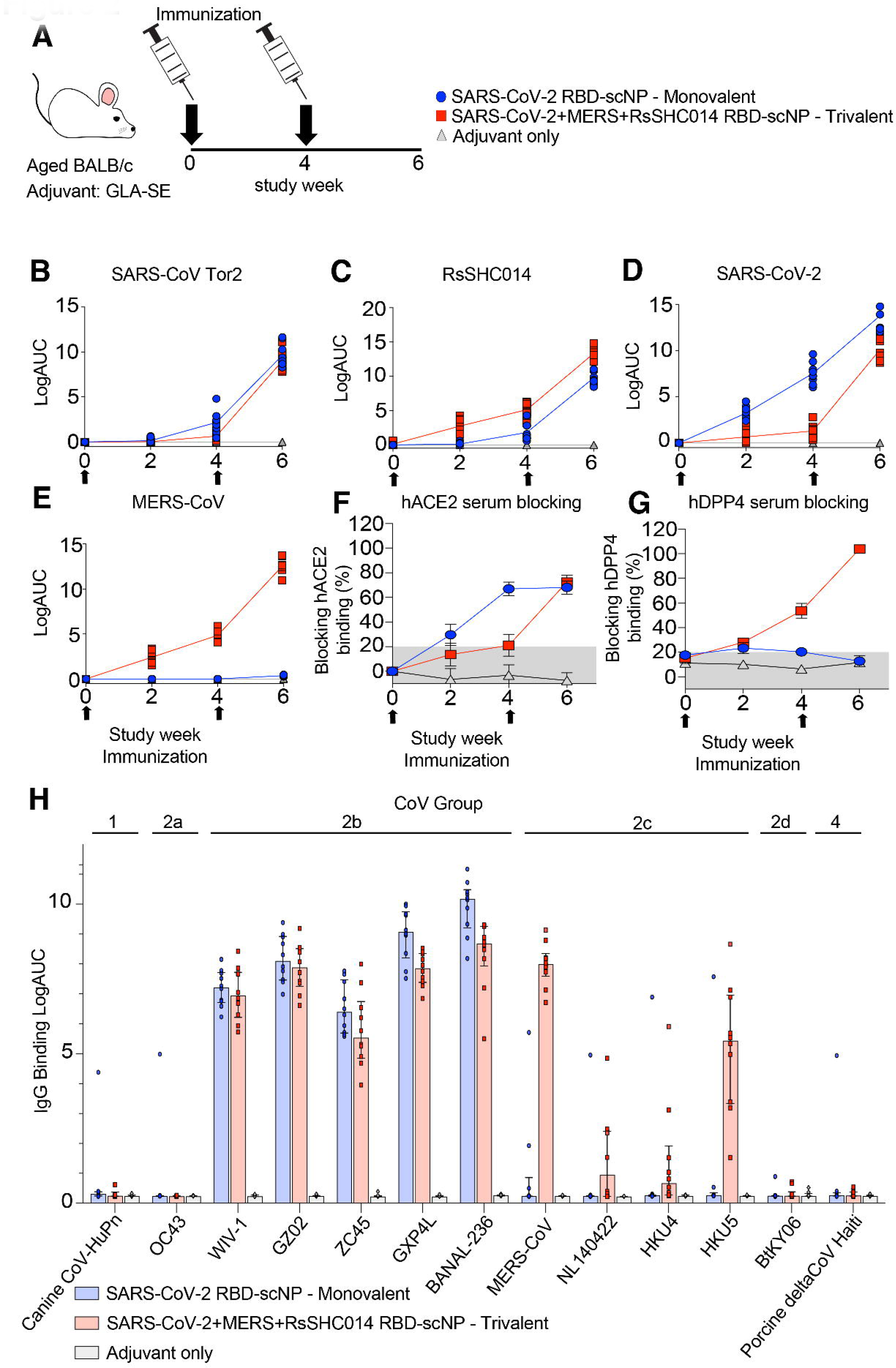
IgG binding responses in mice immunized with monovalent SARS-CoV-2 RBD scNP vaccine, trivalent SARS-CoV-2/RsSHC014/MERS-CoV RBD scNP, and adjuvant alone. Mouse sera was measured at pre-prime, pre-boost, and two-week post boost against the following spike antigens (**A**) SARS-CoV Tor2, (**B**) RsSHC014, (**C**) SARS-CoV-2, and (**D**) MERS-CoV. (**E**) Vaccine-elicited hACE2-blocking serum responses in monovalent, trivalent, and adjuvant-only vaccinated mice. (**F**) Vaccine-elicited hDPP4-blocking serum responses in monovalent, trivalent, and adjuvant-only vaccinated mice. (**G**) Cross-reactivity of monovalent, trivalent, vs adjuvant-only IgG responses against Group 1 (Canine CoV-HuPn), Group 2a (OC43), 2b (WIV-1, SARS-CoV GZ02, ZC45, GXP4L, and BANAL-236), 2c (MERS-CoV, NL140422, HKU4, and HKU5), 2d (BtKY06), and Group 4 (Porcine deltaCoV Haiti) coronavirus RBDs.

To gain insights about the antibody binding breadth and depth elicited by the trivalent RBD scNP vaccine, we assessed serum IgG binding in vaccinated mice to thirteen RBDs of animal and human coronaviruses from Groups 1, 2, and 4 ^2, 3^. The RBD panel included a canine CoV-huPn, OC43, WIV-1, SARS-CoV GZ02, ZC45, GXP4L, BANAL-236, MERS-CoV, NL140422, HKU4, HKU5, BtKY06, and porcine deltacoronavirus Haiti (Figure 2H). The trivalent SARS-CoV-2/RsSHC014/MERS-CoV RBD scNP vaccine elicited high IgG binding responses against five different Group 2b RBDs and four different Group 2c RBDs (Figure 2H). IgG binding was not observed against Group 1, Group 2a, 2d, and Group 4 coronaviruses (Figure 2H). Notably, the MERS-CoV RBD in the trivalent vaccine elicited high binding responses against MERS-CoV and HKU5 and markedly lower binding was observed against NL140422 and HKU4 (Figure 2H). This heterogenous binding across Group 2c RBDs suggests that Group 2c RBDs may share fewer conserved epitopes as compared to Group 2b RBDs (Figure 2H). In contrast, the monovalent SARS-CoV-2 RBD scNP vaccine only elicited high IgG binding responses to Group 2b betacoronaviruses (Figure 2H), demonstrating more limited breadth than the trivalent RBD scNP. Together, these findings indicated that the superior trivalent RBD scNP vaccine elicited broad IgG responses against Group 2b and 2c viruses, and its IgG response exhibited depth reacting with multiple human and animal CoV RBDs from Group 2b and 2c coronaviruses.

### Induction of SARS-CoV-2, RsSHC014, MERS-CoV neutralizing antibodies

We then measured serum neutralizing antibody responses against Group 2b and Group 2c coronaviruses using live-virus assays. At baseline, both the monovalent SARS-CoV-2 RBD and trivalent SARS-CoV-2/RsSHC014/MERS-CoV RBD scNP-vaccinated mice had undetectable neutralizing antibodies against the highly transmissible SARS-CoV-2 BA.1, SARS-CoV Urbani, and MERS-CoV. Following two immunizations, monovalent SARS-CoV-2 RBD scNP vaccinated mice elicited serum neutralizing antibodies against SARS-CoV-2 BA.1 with a median ID_80_ of 1832 (Figure 3A). Similarly, monovalent vaccinated mice elicited potent serum neutralizing antibodies against SARS-CoV Urbani with a median ID_80_ of 1157 (Figure 3B).

**Figure 3.**
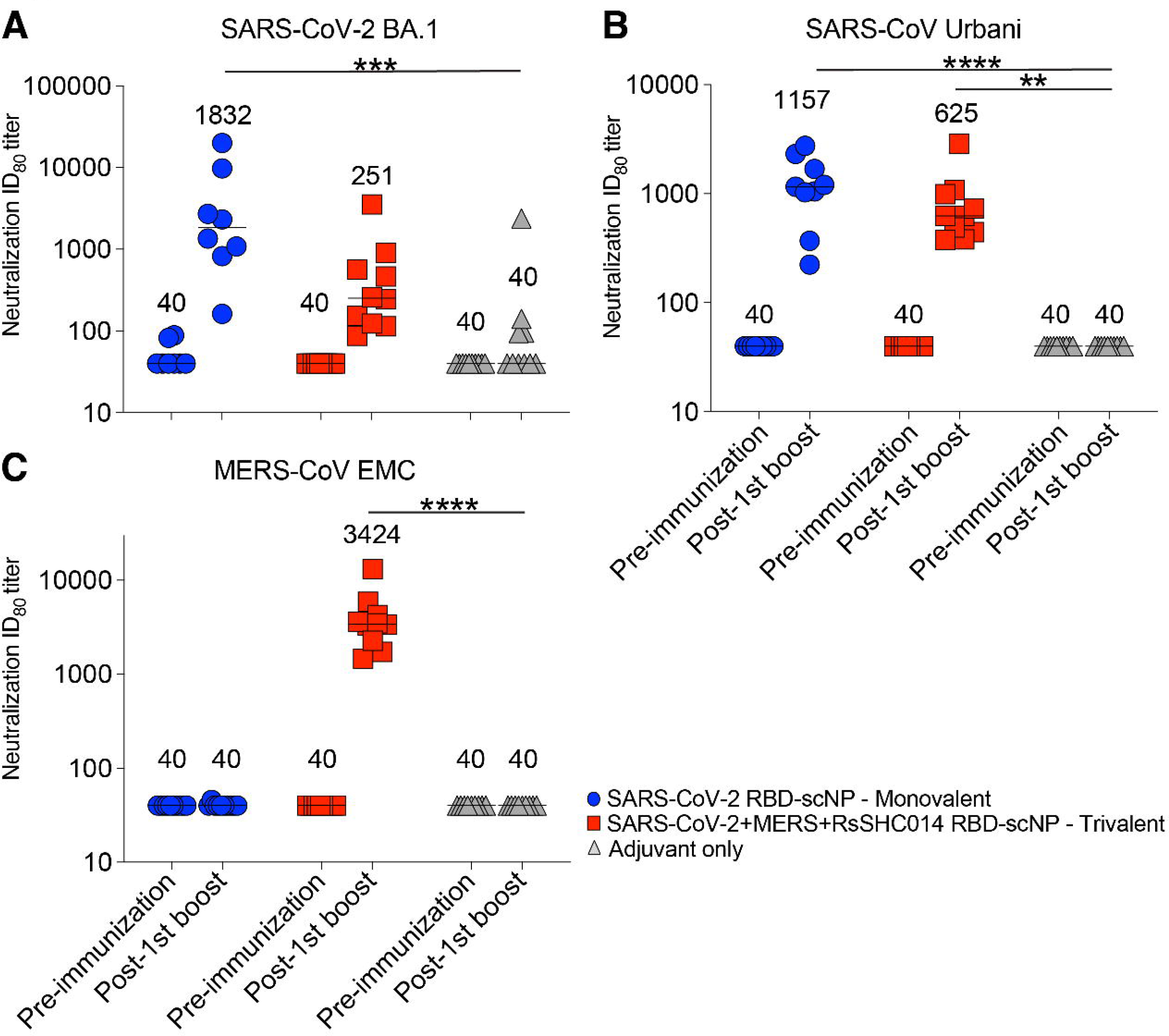
Neutralizing antibodies elicited against Group 2b and Group 2c betacoronaviruses. Live virus neutralizing activity against SARS-CoV-2 BA.1, SARS-CoV Urbani, and MERS-CoV EMC. Mouse sera at baseline and post boost are shown in 2X vaccinated mice are shown. Blue circles denote monovalent SARS-CoV-2 RBD scNP vaccinated mice. Red squares denote trivalent SARS-CoV-2/RsSHC014/MERS-CoV RBD scNP vaccinated mice. Gray triangles denote adjuvant-only control mice. Numerical values in the graphs denote the median ID_80_ values (n = 16; *P < 0.05, **P < 0.005, ***P < 0.0005, and ****P < 0.0001).

Undetectable serum neutralizing antibodies were observed against MERS-CoV EMC (Figure 3C). In contrast to the monovalent vaccine, we observed potent serum neutralizing antibodies against MERS-CoV by the trivalent SARS-CoV-2/RsSHC014/MERS-CoV RBD scNP vaccine with a median ID_80_ of 3424 (Figure 3C). The trivalent SARS-CoV-2/RsSHC014/MERS-CoV RBD scNP vaccine also elicited serum neutralizing antibodies against SARS-CoV-2 BA.1 and SARS-CoV-1 Urbani with ID_80_ values of 251 and 625, respectively. Importantly, undetectable serum neutralizing antibodies were measured in the adjuvant-only vaccinated mice. Thus, the monovalent SARS-CoV-2 RBD scNP vaccine elicited neutralizing antibodies against pandemic and epidemic sarbecoviruses whereas trivalent SARS-CoV-2/RsSHC014/MERS-CoV RBD scNP vaccines elicited neutralizing antibodies against pandemic and epidemic sarbecoviruses and MERS-CoV.

### Protective efficacy of trivalent RBD nanoparticle vaccine against Group 2b and Group 2c CoVs

To evaluate the protective efficacy of the trivalent RBD scNP against sarbecovirus and merbecovirus infection with highly pathogenic coronaviruses, we challenged mice with either a heterologous, lethal mouse-adapted SARS-CoV virus (MA15) ^22^, or a highly pathogenic mouse-adapted MERS-CoV virus (m35c4) ^23, 24^. Aged BALB/c mice immunized with the trivalent SARS-CoV-2/RsSHC014/MERS-CoV RBD were protected from weight loss (Figure 4A) and mortality (Figure 4B) after SARS-CoV MA15 challenge. This protection was likely due to conserved RBD epitopes shared among sarbecoviruses ^7, 10, 11, 25^. Notably, the monovalent SARS-CoV-2 RBD scNP vaccine also protected against heterologous SARS-CoV MA15 challenge, whereas the adjuvant-only-vaccinated controls had 40% mortality by day 4 post infection (Figure 4B). Compared to adjuvant-only controls, both monovalent and trivalent RBD scNP had reduced lung virus replication at day 2 post infection as measured by infectious virus plaque assays (Figure 4C). However, only the trivalent scNP-vaccinated mice had lower infectious SARS-CoV replication in the nasal turbinates at day 2 post infection as measured by plaque assay compared to the adjuvant-only-vaccinated controls (Figure 4D). Moreover, the trivalent RBD scNP vaccine also mediated increased protection against upper airway replication of SARS-CoV in mice.

**Figure 4.**
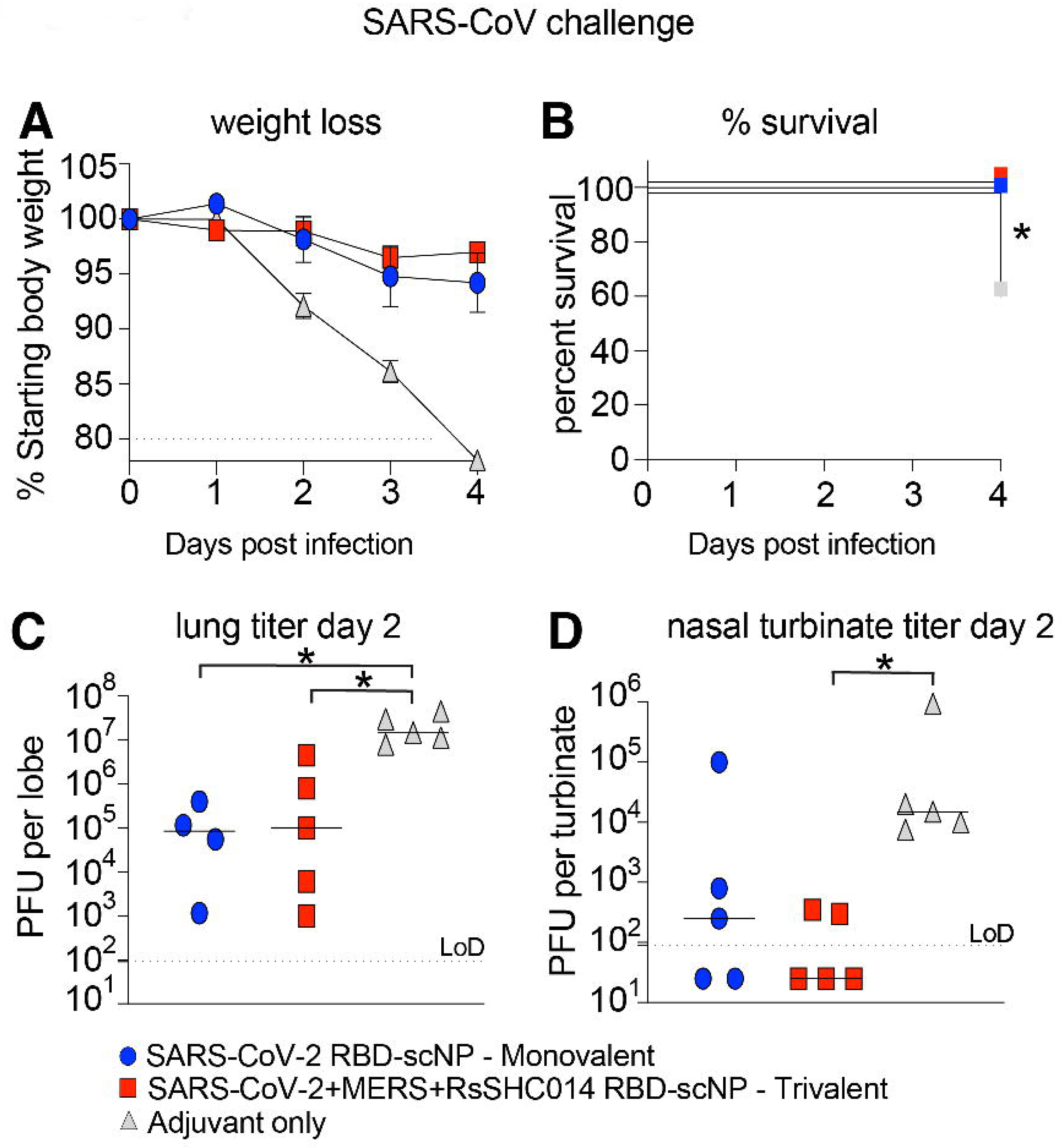
Protective efficacy of monovalent vs trivalent RBD scNP vaccines against SARS-CoV challenge in mice. (**A**) Weight loss in monovalent SARS-CoV-2 RBD, trivalent SARS-CoV-2/RsSHC014/MERS-CoV RBD vaccinated scNP, and adjuvant-only vaccinated mice. (**B**) Percent survival in vaccinated mice vs control following lethal SARS-CoV Urbani MA15 challenge. Statistical significance of the survival curves is from a Chi square log-rank test. (**C**) Infectious virus replication in the lung of vaccinated mice at day 2 following infection. Statistical significance is from a Kruskal-Wallis test following a Dunn’s multiple comparison correction test. (**D**) Infectious virus replication in nasal turbinates at day 2 post infection. Statistical significance is from a Kruskal-Wallis test following a Dunn’s multiple comparison correction test. Blue circles represent the monovalent vaccinated mice. Red squares represent the trivalent vaccinated mice. Grey triangles denote the adjuvant-only vaccinated mice. *P < 0.05, **P < 0.005, ***P < 0.0005, and ****P < 0.0001

As we observed strong protection from heterologous and highly pathogenic SARS-CoV MA15, we evaluated whether the trivalent vaccine also protected against challenge in a highly pathogenic mouse-adapted MERS-CoV model ^23, 24^. Like adjuvant-only controls, DPP4 transgenic mice vaccinated twice with a monovalent SARS-CoV-2 RBD scNP vaccine experienced severe MERS-CoV disease including weight loss (Figure S2A), high levels of infectious virus replication in the lung and nasal turbinates (Figure S2B and C). Similarly, by day 4 post infection SARS-CoV-2 RBD scNP-vaccinated mice exhibited significant weight loss and high amounts of virus replication in the lung (Figure S2D). In contrast, mice vaccinated twice with the trivalent SARS-CoV-2/RsSHC014/MERS-CoV RBD scNP vaccine were protected from weight loss (Figure S2A). Unlike adjuvant-only controls and SARS-CoV-2 RBD monovalent vaccinated mice, we observed complete protection from lung virus replication at day 2 post infection in the trivalent SARS-CoV-2/RsSHC014/MERS-CoV RBD scNP 2X vaccinated group (Figure S2B). However, we did not observe complete suppression of nasal turbinate MERS-CoV replication at day 2 post infection and lung virus replication at day 4 post infection in the trivalent SARS-CoV-2/RsSHC014/MERS-CoV RBD scNP 2X vaccinated group (Figure S2C-D).

To evaluate if additional boosting could increase the protective efficacy in the upper airways of the trivalent SARS-CoV-2/RsSHC014/MERS-CoV RBD scNP vaccine, we repeated the vaccination study in the DPP4-modified mice that are susceptible to MERS-CoV infection and disease. We vaccinated mice three times four weeks apart with either the trivalent RBD scNP or SARS-CoV-2 RBD scNP (Figure S3A). Notably, mice immunized three times (3X) showed an increase in serum binding IgG against Group 2b and 2c coronavirus RBDs indicating the additional boost augmented antibody responses (Figure S3B). Three immunizations with the trivalent SARS-CoV-2/RsSHC014/MERS-CoV RBD scNP vaccine completely protected mice from weight loss (Figure 5A), and lung virus replication at days 3 and 5 following MERS-CoV challenge (Figure 5B, 5D). Importantly, mice vaccinated 3x with the trivalent vaccine were fully protected from MERS-CoV replication in the nasal turbinates (Figure 5C). In contrast, the adjuvant-only control and the SARS-CoV-2 RBD monovalent scNP vaccine group exhibited marked weight loss following MERS-CoV challenge (Figure 5A) and had high levels of virus replication in the lungs and nasal turbinates at day 3 (Figure 5B and C). At day 5 post infection, virus replication remained high in these two groups of mice (Figure 5D). Therefore, a three-dose vaccination strategy achieved a high degree of protection in both the lower and upper airway after challenge with MERS-CoV.

**Figure 5.**
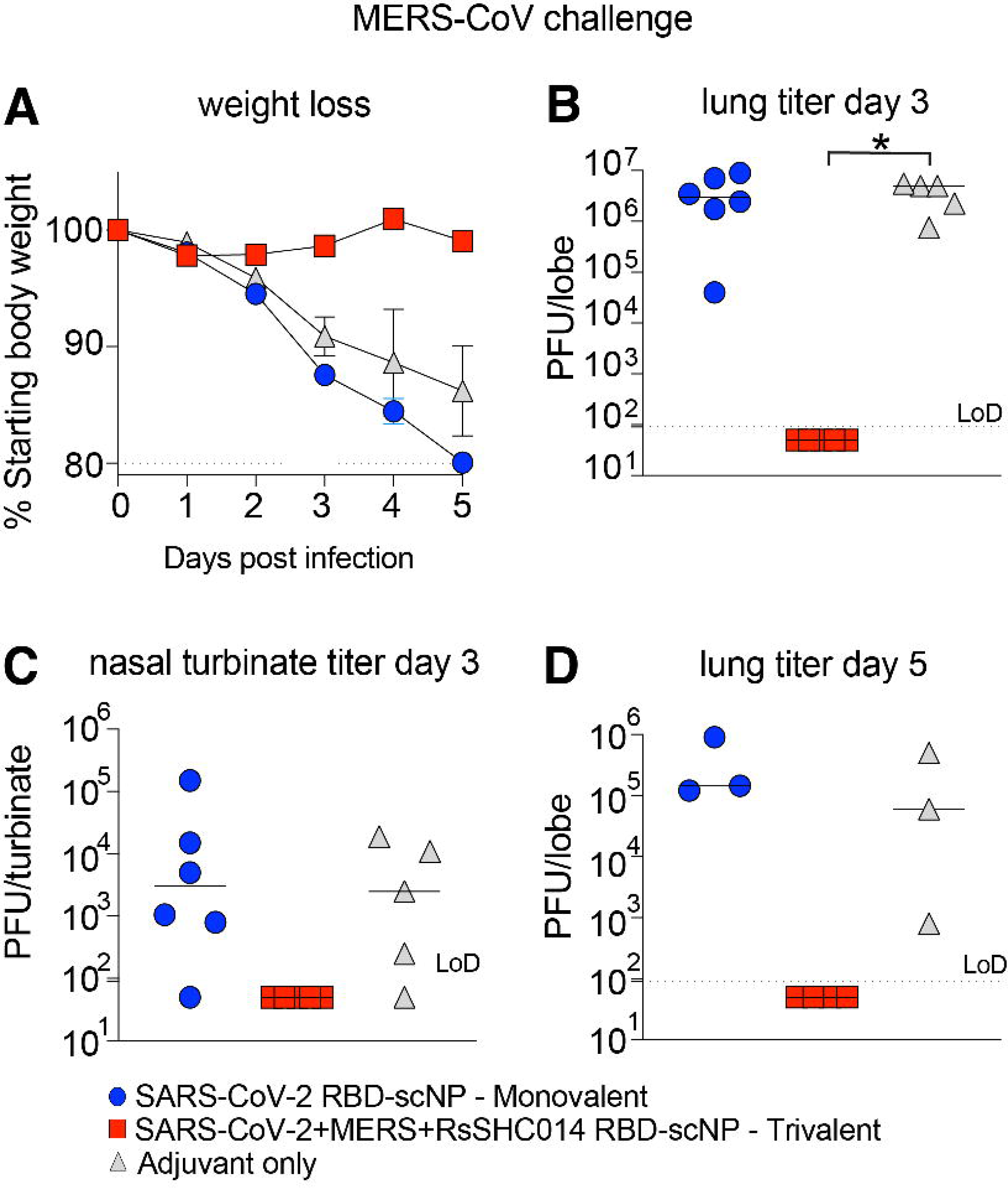
Protective efficacy of monovalent vs trivalent RBD scNP vaccines against MERS-CoV challenge in mice. (**A**) Weight loss in SARS-CoV-2 RBD monovalent, SARS-CoV-2/RsSHC014/MERS-CoV RBD vaccinated scNP, and adjuvant-only vaccinated mice following MERS-CoV intranasal challenge. Lung virus replication in monovalent, trivalent, and adjuvant-only controls at day 3 post infection. (**C**) Infectious virus replication in nasal turbinates at day 3 post infection. (**D**) Lung infectious virus replication at day 5 post infection. P values shown in all panels are from a Kruskal-Wallis test following a Dunn’s multiple comparisons test. *P < 0.05, **P < 0.005, ***P < 0.0005, and ****P < 0.0001

## DISCUSSION

Given the more than 6.8 million deaths attributed to the SARS-CoV-2 pandemic, vaccines that protect against the known highly pathogenic human coronaviruses are needed ^26, 27^. This current study demonstrated a trivalent receptor binding domain sortase-conjugated nanoparticle vaccine induced neutralizing antibodies against all three highly pathogenic human betacoronaviruses and protected against both heterologous Group 2b (*Sarbecovirus* subgenus) and homologous Group 2c (*Merbecovirus* subgenus) coronavirus infections. This vaccine is an advance over current SARS-CoV-2 mRNA vaccines, which lack protection against other human pathogenic betacoronaviruses such as SARS-CoV and MERS-CoV ^6^. The trivalent vaccine is also an advance beyond current Group 2b-focused RBD nanoparticle vaccines. The monovalent SARS-CoV-2 RBD scNP vaccine used in this study elicited high concentrations of IgG antibodies against Group 2b RBDs, and in previous studies was shown to neutralize recent known SARS-CoV-2 variants including highly mutated BA.4/BA.5 omicron sub-strains ^10^.

Moreover, the monovalent SARS-CoV-2 RBD scNP vaccine protects against sarbecoviruses SARS-CoV, SARS-CoV-2, and RsSHC014 ^10^. However, this SARS-CoV-2 RBD nanoparticle did not generate cross-reactive antibodies against Group 2c spike ^7^. Notably, monovalent scNP SARS-CoV-2 vaccines that protect mice and monkeys against SARS-CoV-2 and sarbecovirus challenge ^7, 10^ did not protect against MERS-CoV challenge. The lack of broadly reactive Group 2b and 2c antibodies is expected given that MERS-CoV and SARS-CoV-2 RBDs differ in overall structure ^28, 29^. Therefore, “universal” vaccine approaches targeting SARS-CoV-2 variants may be distinct from those approaches needed for vaccines against antigenically and genetically distant coronaviruses.

Importantly, the SARS-CoV-2 RBD was sufficient in the monovalent vaccine for eliciting cross-reactive IgG antibodies against all tested sarbecoviruses. To bolster immunity against sarbecoviruses, the trivalent RBD nanoparticle includes SHC014 RBD. Conversely, the MERS-CoV RBD in the trivalent RBD vaccine elicited a range of high and low binding IgG titers to the four Group 2c RBDs tested. The inability of a single Group 2c RBD to elicit high titers of cross-reactive IgG to all Group 2c RBDs tested indicates that Group 2c RBDs may share less epitope conservation compared to Group 2b RBDs.

The development of antibody-based MERS vaccines has been of concern given reports that antibody dependent enhancement of infection can occur *in vitro* with MERS-CoV-reactive antibodies ^30^. Increased virus replication that is mediated by IgG antibodies is a classical surrogate of antibody-dependent enhancement that is observed for flaviviruses like dengue virus ^31^. In our study, we observed potent serum antibody neutralization of MERS-CoV *in vitro* and no evidence of increased virus replication upon challenge of mice immunized with MERS-CoV RBD. It is also important to note that we did not observe increased lung or nasal turbinate MERS-CoV replication relative to adjuvant only controls in mice vaccinated with the monovalent SARS-CoV-2 RBD scNP vaccine even though this vaccine did not protect against MERS-CoV challenge. This is an important observation as it suggests that individuals that have SARS-CoV-2 immunity to the RBD are unlikely to experience more severe disease when exposed to MERS-CoV or to a distinct Group 2c coronavirus that is antigenically like MERS-CoV.

A limitation to our study is that mucosal antibody responses were not measured. Thus, the durability of vaccine-elicited neutralizing antibodies in the upper airway is not known. Similarly, tissue-resident memory B and T cells responses were not profiled in the nasal airways or lungs ^17^. Another limitation to our study is the lack of a heterologous Group 2c challenge in the trivalent scNP vaccine expressing MERS-CoV RBD. However, this is currently a limitation of the broad coronavirus pathogenesis field as MERS-CoV is the only known Group 2c human respiratory coronavirus that can replicate and cause disease in mice expressing humanized DPP4 receptors.

Finally, our study shows the utility of the sortase conjugate nanoparticle platform for rapidly and easily generating broadly protective vaccines. The trivalent RBD scNP vaccine is a viable strategy for vaccine-mediated protection against the three highly pathogenic Group 2b and 2c betacoronaviruses - SARS-CoV, SARS-CoV-2 and its variants, and MERS-CoV. Moving forward it will be critical to assess if this trivalent RBD scNP vaccine also protect against Group 2b and Group 2c coronaviruses in additional mouse models that express human ACE2 in the upper and lower airway epithelium as is observed in humans ^32^ and in other MERS-CoV mouse challenge models ^33^. Our results suggest that universal vaccine approaches targeting Group 2b and Group 2c coronaviruses is achievable via multivalent delivery of RBDs via adjuvanted nanoparticle vaccines. The protective Group 2c immunity generated by the trivalent RBD nanoparticle is important since previous MERS-CoV outbreaks have had case fatality rates as high as 40% ^34^, far exceeding the 1-10% rate reported for SARS-CoV-2 and SARS-CoV ^35^. The next generation of coronavirus vaccines will need to broaden protection to include both Group 2b and 2c coronaviruses. Additionally, these findings have important implications for slowing down or preventing the spread of pre-emergent, zoonotic coronaviruses poised for human emergence^19, 20^_._

## STAR METHODS

### RBD sortase A conjugated nanoparticle vaccine production

The receptor-binding domains (RBDs) from SARS-CoV-2 Wuhan-Hu1 isolate, MERS-CoV EMC isolate, and BatCoV RsSHC014 were expressed with a Sortase A donor sequence (LPETGG) encoded at the C terminus. An HRV-3C cleave site, an 8x His-tag, and a twin StrepTagII (IBA) were added C-terminal to the Sortase A donor sequence. The RBDs were each expressed by transient transfection using 293Fectin in Freestyle 293 cells and purified by StrepTactin affinity chromatography (IBA) followed by Superdex200 size-exclusion chromatography as described previously^7, 36^. *Helicobacter pylori* ferritin particles were expressed with an N-terminal pentaglycine Sortase A acceptor sequence at the end of each subunit. 6x His-tags were included C-terminal to an HRV-3C cleavage site to enable affinity purification of the Ferritin particles. Prior to conjugation, RBDs, ferritin subunits, and pentamutant Sortase A ^37^ were buffer exchanged into 50mM Tris, 150mM NaCl, 5mM CaCl_2_ at pH 7.4. The components were combined at a ratio of 360 μM total RBD (360 μM SARS-CoV-2 RBD for monovalent RBD scNP, or 120μM each of SARS-CoV-2, RsSHC014, and MERS-CoV RBD for the trivalent RBD scNP), plus 120uM Ferritin, plus 100uM Sortase A, and incubated at room temperature for 4 hours. After incubation, the conjugated RBD-bearing nanoparticles were separated from free unconjugated reactants by size-exclusion chromatography using a Superose6 16/600 column. Conjugate nanoparticle assembly was confirmed by NSEM and by Western blot under both reducing and non-reducing conditions.

### Biolayer interferometry (BLI)

Antibody binding was determined using a FortéBio Bio-Layer Interferometry instrument (Sartorius Octet Red96e) at 25°C with a shake speed of 1000 rpm. Antibodies were diluted to 20 μg/mL in a flat bottom 96-well plate (Greiner) with .22μm filtered phosphate buffered saline pH 7.4 and 0.05% Tween 20 (PBS-T). The antigens were diluted to a concentration of 50μg/mL using PBS-T. Hydrated Anti-hIgG Fc Capture (AHC) biosensors (Sartorius #18-5060) were equilibrated for 60 second and then antibodies were loaded to biosensors for 300 seconds. After a 60-second wash and a 180-second baseline step, biosensors were then dipped into the diluted antigens for a 200-second association. Next, antibody and antigens allowed to dissociate for 300 seconds. Data was analyzed using Data Analysis HT 12.0 software. The negative control antibody, CH65, was indicated as a reference sensor and subtracted from the remaining ligand sensor measurements. Data was then aligned to the average of the baseline step and plotted using GraphPad Prism 9 software.

### Negative stain electron microscopy of RBD nanoparticles

Negative stain electron microscopy was performed as previously described ^7^. The RBD nanoparticle protein was thawed in an aluminum block at room temperature for 5 min. The RBD scNP was diluted to a final concentration of 0.2 mg/mL into room-temperature buffer containing 150 mM NaCl, 20 mM HEPES pH 7.4, 5g/dL glycerol and 8 mM glutaraldehyde. After 5 min, the cross-linking was quenched by the addition of 1 M Tris pH 7.4 to a final concentration of 75 mM Tris and incubated for 5 min. Carbon-coated grids (EMS, CF300-cu-UL) were glow-discharged for 20 s at 15 mA, and subsequently a 5-μl drop of quenched sample was incubated on the grid for 10–15 s. The grid was blotted and then stained with 2g/dL uranyl formate. After air drying, grids were imaged with a Philips EM420 electron microscope operated at 120 kV, at 49,000× magnification and images captured with a 76 megapixel CCD camera at a pixel size of 2.4 Å.

### Processing of negative-stain images

The RELION 3.0 program was used for all negative-stain image processing following previously published procedures ^7^. Images were CTF-corrected with CTFFIND and particles were picked using a nanoparticle template. Extracted particle stacks were underwent 2 or 3 rounds of 2D class averaging and selection to discard irrelevant particles and background picks.

### Mouse vaccinations and virus challenge experiments

Aged BALB/c (#047) retired breeder female mice were purchased from Envigo and were used for SARS-CoV-1 MA15 challenge studies. B6 male and female mice modified at the DPP4 locus ^23^ to allow pathogenesis by mouse-adapted MERS-CoV m35c4 ^24^were bred in house and used at ∼20-25 weeks of age. The Toll-like receptor 4 agonist glucopyranosyl lipid adjuvant–stable emulsion (GLA-SE) was used as the adjuvant for the vaccine immunogens. Mouse vaccination studies were performed intramuscularly with GLA-SE-adjuvanted SARS-CoV-2 RBD scNP, GLA-SE-adjuvanted SARS-CoV-2/RsSHC014/MERS-CoV RBD scNP, or GLA-SE-adjuvant only for the control group. Vaccine immunogens were administered at 10 μg of the RBD scNP vaccines formulated with 5 μg of adjuvant. Mice were immunized at week 0 and week 4 for the 2X prime-boost vaccine regimen, and at week 0, week 4, and week 8 for the 3X prime-boost-boost vaccine regimen. Mice were then moved into the BSL3 and acclimated for a few days.

Prior to challenge, mice were anesthetized with an intraperitoneal delivery of xylazine and ketamine and given a lethal dose of SARS-CoV-1 MA15 ^22^: 1L×L10^4^ PFU/ml. For the MERS-CoV challenge studies, mice were challenged with mouse-adapted MERS-CoV m35c4 ^24^.

### Binding ELISA against coronavirus antigen panel

For coronavirus antigen-binding assays, 384-well ELISA plates (Costar #3700) were coated with 2 μg/ml antigens in 0.1M sodium bicarbonate overnight at 4°C. Plates were then washed 1X and blocked for 2 h at room temperature with SuperBlock (1X phosphate buffered saline (PBS) containing 4% (w/v) whey protein 15% normal goat serum/0.5% Tween-20/0.05% sodium azide). Mouse serum samples were collected at baseline before prime, two weeks post prime, four weeks post prime, two weeks post boost, and two weeks post the second boost. Mouse serum samples were added at 1:30 dilution in SuperBlock and diluted 3-fold through 12 dilution spots to generate binding curves. Diluted serum samples were bound to coated plates in SuperBlock for 1h at room temperature. Plates were then washed 2X and a horseradish peroxidase (HRP)-conjugated goat anti-mouse IgG secondary antibody (SouthernBiotech 1030-05) was added in SuperBlock at a 1:16,000 dilution. Secondary antibody was bound for 1h and then washed 4X and detected with 20μL SureBlue Reserve (KPL 53-00-03) for 15 min.

Colorimetric reactions were stopped by adding 20μL of 1% HCL stop solution. Plates were read at 450nm and area under the curve (AUC) was calculated from the serially diluted mouse serum samples.

### Live virus neutralization assays

All live virus assays were performed in a BSL-3 laboratory. Full length SARS-CoV Urbani, SARS-CoV-2 Wuhan-1 expressing the BA.1 spike, and MERS-CoV were designed to express nanoluciferase (nLuc) as described previously ^38, 39^. SARS-CoV Urbani and SARS-CoV-2 BA.1 stocks were generated and titrated in Vero E6 (C1008) cells and MERS-CoV stocks were titrated in Vero 81 (CCL-81) cells. For the live virus neutralization assays, cells were plated at 20,000 cells per well in clear bottom, black-walled 96-well plates the day prior to the assay. On the day of the assay, mouse serum samples diluted 1:40 and serially diluted 3-fold to eight dilutions.

Serially diluted mouse serum was added at a 1:1 volume with diluted virus and incubated for 1 h. Antibody-virus dilutions were then added to cells at 800 PFU per well and incubated at 37°C with 5% CO_2_. Following a 24hr incubation, plates were read by adding 25μL of Nano-Glo Luciferase Assay System (Promega). Luminescence was measured by a Spectramax M3 plate reader (Molecular Devices). Fifty percent virus neutralization titers were calculated using GraphPad Prism via four-parameter dose-response curves.

### Biocontainment and biosafety

All experiments handling live viruses, including mouse-adapted coronaviruses, were performed in an animal biosafety level 3 (BSL-3) laboratory. Laboratory workers performing BSL-3 experiments wore powered air purifying respirators (PAPR), Tyvek coverall suits, double booties covering footwear, and double gloves. All recombinant coronavirus work was approved by the UNC Institutional Biosafety Committee (IBC). All animal work was approved by the UNC Institutional Animal Care and Use Committee (IACUC). All BSL-3 work was performed in a facility conforming to requirements recommended in the Microbiological and Biomedical Laboratories, by the U.S. Department of Health and Human Services, the U.S. Public Health Service, and the U.S. Center for Disease Control and Prevention (CDC), and the National Institutes of Health (NIH).

### Statistical analysis

Non-parametric Kruskal-Wallis tests were used to compare lung and nasal turbinate infectious virus replication in challenged mice and neutralizing antibody assays. A Dunn’s test was used to correct for multiple comparisons. A Chi square log-rank test was used for the survival analysis. Statistical analyses were performed in GraphPad Prism 9.

## Funding

D.R.M. is supported by a Hanna H. Gray Fellowship from the Howard Hughes Medical Institute. This project was supported by the NIAID, NIH, U.S. Department of Health and Human Services award U54 CA260543 (to R.S.B.), and AI158571 (to B.F.H.), as well as an animal models contract from the NIH (HHSN272201700036I).

## Author contributions

D.R.M., R.S.B. B.F.H. and K.O.S. conceived the study. D.R.M., T.D.G., A.S., R.S.B., B.F.H., K.O.S., designed experiments. D.R.M., T.D.G., A.S., M.L.M, N.J.C., K.G., T.S., R.J.E., K.M.,performed laboratory experiments. D.R.M., T.D.G., A.S., M.L.M, N.J.C., K.G., T.S., R.S.B., B.F.H., K.O.S. analyzed the data and provided critical insight. D.R.M. wrote the first draft of the paper. D.R.M., T.D.G., A.S., M.L.M, N.J.C., K.G., T.S., R.S.B., B.F.H., K.O.S. edited the paper. All authors read and approved the final version of the paper.

## Competing interests

B.F.H. and K.O.S. have filed US patents regarding the nanoparticle vaccine. R.S.B. is on the scientific advisory boards of VaxArt, Invivyd, and Takeda.

## Data and materials availability

All data associated with this paper is in the paper and Supplementary Materials. Reagents developed in this study, including vaccine immunogens and viruses will be made available by contacting R.S.B., B.F.H. and K.O.S. following the completion of a material transfer agreement.

---

**Figure S1.**
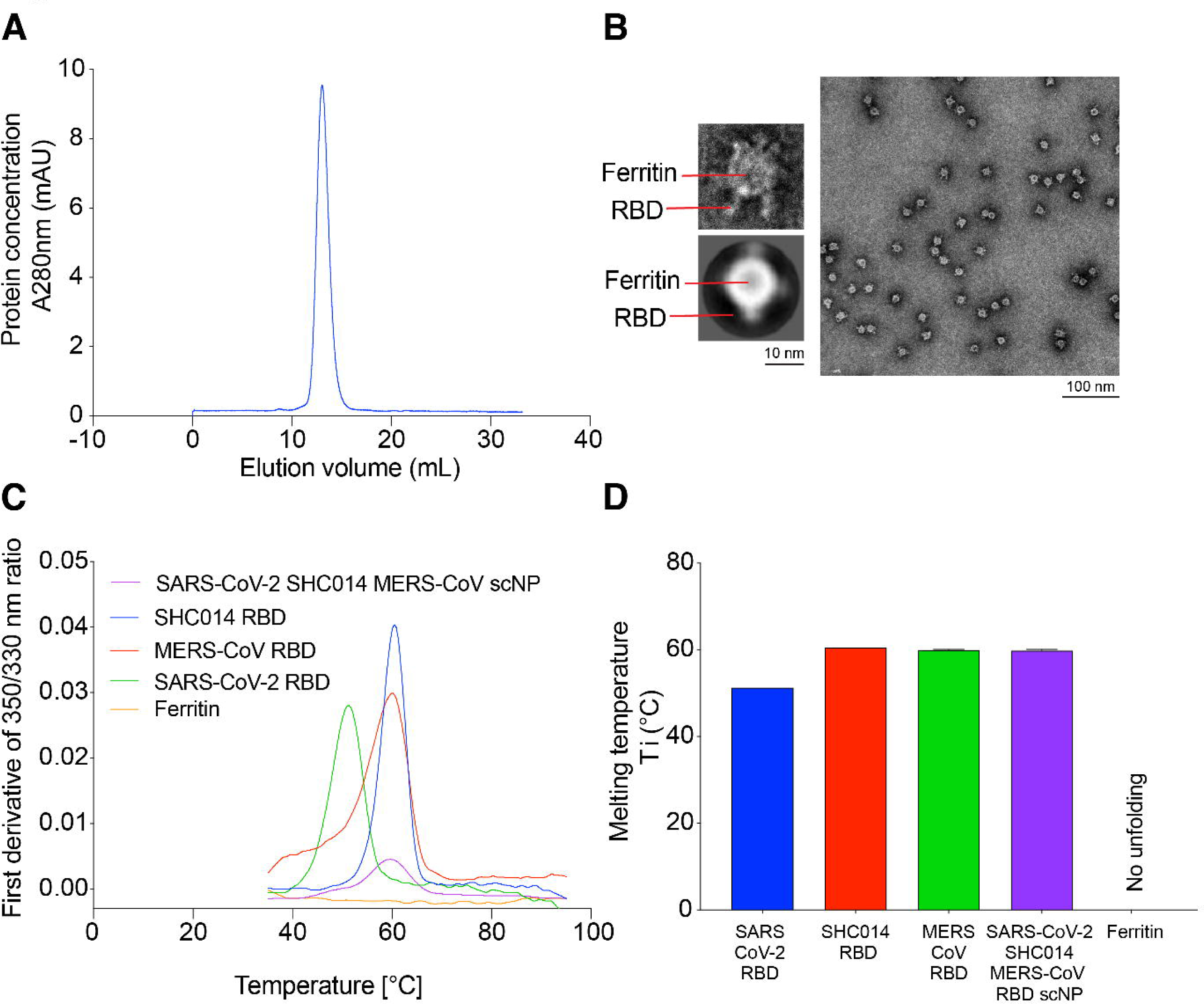
Structural and biophysical characterization of the trivalent SARS-CoV-2/SHC014/MERS RBD scNP. **(A)** Analytical size exclusion chromatography with a Superose6 column showing a homogenous protein nanoparticle at the expected elution volume. **(B)** Negative stain electron microscopy of trivalent RBD scNP. 2D class average is shown on the bottom left and a raw image of a single nanoparticle is shown on the top left. The raw image of the carbon grid is shown on the right. **(C and D)** Differential scanning fluorimetry of the individual components and assembled trivalent RBD scNP. Melting temperature is defined as the inflection temperature (T_i_) on the 350 nm/330 nm ratio curve.

**Figure S2.**
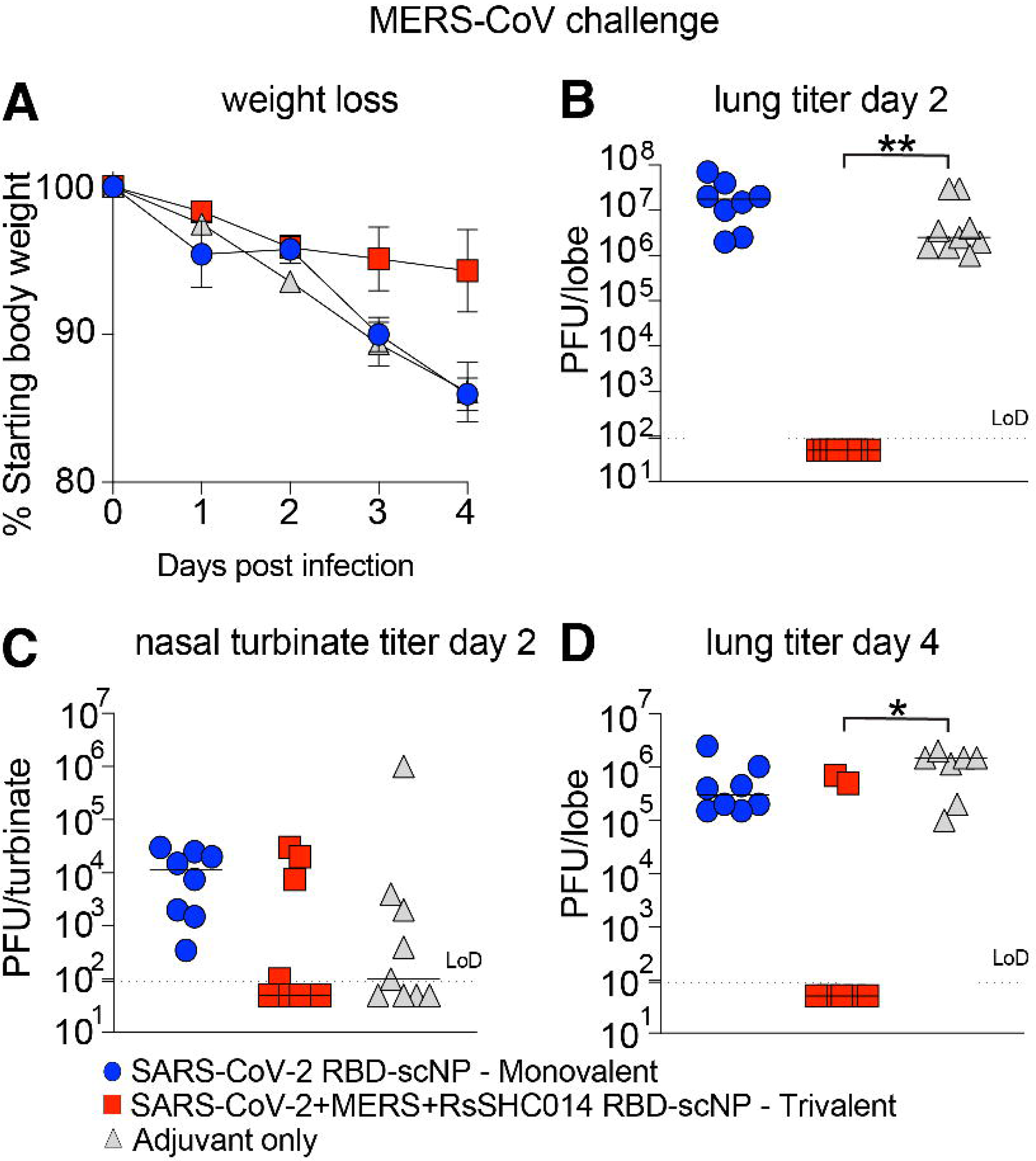
Protective efficacy of monovalent vs trivalent RBD scNP vaccines against MERS-CoV challenge in 2X vaccinated mice. (**A**) Weight loss following MERS-CoV intranasal challenge in monovalent, trivalent, and adjuvant-only vaccinated mice. (**B**) Infectious virus replication in the lung at day 2 post infection. (**C**) Infectious virus replication at day 2 post infection in nasal turbinates. (**D**) Infectious virus replication at day 4 post infection. A Kruskal-Wallis test with a Dunn’s multiple comparison correction test was used for calculating statistical significance in all panels. *P < 0.05, **P < 0.005, ***P < 0.0005, and ****P < 0.0001

**Figure S3.**
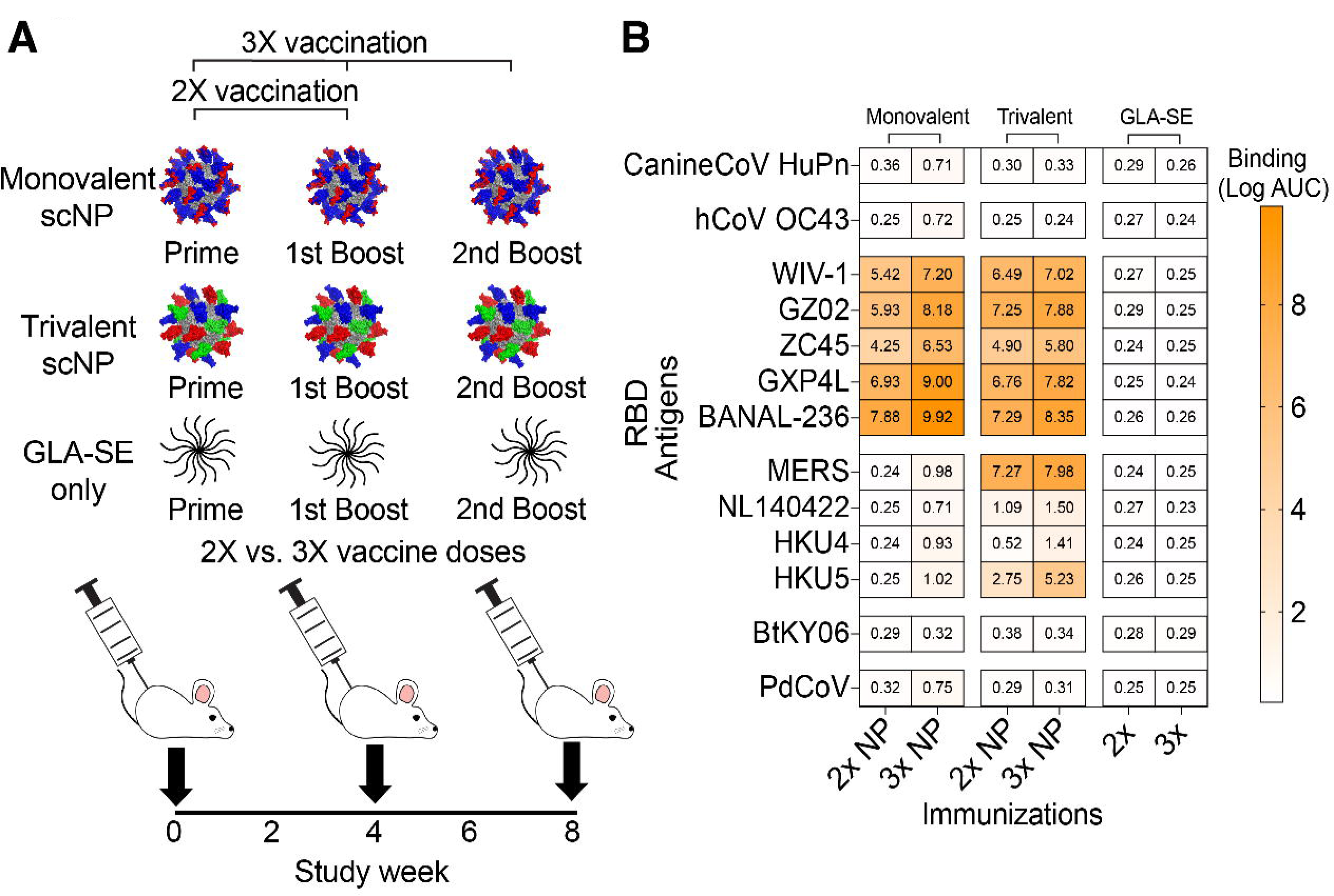
Immunogenicity of monovalent vs trivalent RBD scNP vaccines in 2X vs 3X vaccinated mice. (**A**) vaccination schema with monovalent scNP, trivalent scNP, and adjuvant only (GLA-SE). (**B**) LogAUC IgG binding comparison of 2X (prime-boost) vs 3X (prime-boost-boost) vaccinated mice with monovalent, trivalent, and adjuvant-only against genetically divergent RBDs from Group 1 (canineCoV-HuPn). Group 2a (OC43), Group 2b (WIV-1, GZ02, ZC45, GXP4L, and BANAL-236), Group 2c (MERS-CoV, NL140422, HKU4, and HKU5), Group 2d (BtKY06), and Group 4 (Porcine deltacoronavirus Haiti).

